# GeoWaVe: Geometric median clustering with weighted voting for ensemble clustering of cytometry data

**DOI:** 10.1101/2022.06.30.496829

**Authors:** Ross J. Burton, Simone M. Cuff, Matt P. Morgan, Andreas Artemiou, Matthias Eberl

**Author notes:** The authors wish it to be known that, in their opinion, the last two authors should be regarded as Joint Senior Authors.

## Abstract

**Motivation:** Clustering is an unsupervised method for identifying structure in unlabelled data. In the context of cytometry, is typically used to categorise cells into subpopulations of similar phenotype. However, clustering is greatly dependent on hyperparameters and the data to which it is applied as each algorithm makes different assumptions and generates a different ‘view’ of the dataset. As such, the choice of clustering algorithm can significantly influence results, and there is often not one preferred method but different insights to be obtained from different methods. To overcome these limitations, consensus approaches are needed that directly address the effect of competing algorithms, which to our knowledge has not been applied to cytometry.

**Results:** We present a novel ensemble clustering methodology based on geometric median clustering with weighted voting (GeoWaVe). Compared to graph ensemble clustering methods that have gained popularity in scRNA-seq analysis, GeoWaVe performed favourably on different sets of high-dimensional mass and flow cytometry data. Our findings provide proof of concept for the power of consensus methods to make the analysis, visualisation and interpretation of cytometry data more robust and reproducible. The wide availability of ensemble clustering methods is likely to have a profound impact on our understanding of cellular responses, clinical conditions, and therapeutic and diagnostic options.

**Availability and implementation:** GeoWaVe is available as part of the CytoCluster package https://github.com/burtonrj/CytoCluster.

## 1 Introduction

Clustering is an unsupervised method for identifying structure in unlabelled data. In the context of cytometry, the objective is to categorise events into groups of similar phenotype. This technique is increasingly being adopted in the field and is widely regarded as an acceptable alternative to manual analysis [1-3]. However, the choice of algorithm appears to be often driven either by its availability in commercial software or ease of its use. In many instances the reason behind the particular choice of algorithm is not discussed at all. Of note, clustering algorithms differ in the assumptions made of data, their performance tends to be highly data-specific and results can vary widely depending on the chosen hyperparameters [4-6].

Ensemble clustering (also referred to as consensus clustering) offers an opportunity to reduce this frequently encountered bias by combining the partitions of multiple clustering algorithms run on the same data to identify a consensus that is informed by multiple ’views’, thereby reducing the dependence on any individual algorithm. Unlike ensemble methods in supervised classification, ensemble clustering has many challenges: the number of clusters may differ amongst the base partitions, the optimal number of consensus clusters is often unknown, and it is necessary to solve the correspondence issue of matching clusters between individual partitions [4, 7].

Broadly speaking, ensemble clustering methods can be grouped into three categories: co-association methods, feature-based methods and methods using graph representations [4, 7, 8].

Co-association methods act on the pairwise similarity of clusters sourced from different algorithms. Consensus solutions can be derived from simple techniques such as agglomerative clustering of the binary co-association matrix (*N* × *N* matrix, where *N* is the number of events, for instance the number of single cells) [5] or the cluster-based similarity partitioning algorithm (CSPA), that forms partitions on the derived similarity graph using the METIS software [9].

Methods that act on co-association are burdened by space complexity and are therefore intractable for large data where such a matrix exceeds the available computer memory [4].

Feature-based methods offer an alternative by presenting the problem as a label-association matrix (m × n matrix, where m is the number of unique clusters). Consensus solutions can be formulated with iterative voting, finite mixture models, pairwise agreement between clusters, or agglomerative clustering of this label-association matrix [7].

Another popular approach for consensus clustering is by using graph-based methods, where a weighted graph of the clusters contributing to an ensemble is generated and then partitioned into *k* parts using a graph partitioning technique [4, 7]. Strehl and Ghosh [9] developed the hyper-graph partitioning algorithm (HGPA) and the meta-clustering algorithm (MCLA), both heuristics that represent the clustering ensemble as a hypergraph. Later the hybrid bipartite graph formulation (HBGF) algorithm was introduced as an alternative approach that models clusters and observations in the same graph. In each case, consensus partitions are constructed from a subsequent bipartite graph [10]. The advantage of the aforementioned graph methods is their heuristic approach that avoids the need for a co-association matrix, making them applicable to large data.

Ensemble clustering methods have successfully been adopted in the field of single-cell RNA sequencing (scRNA-seq) but the methodologies chosen usually reflect the size of data generated by this technique and do not address the space complexity issues that arise from larger datasets. the graph partitioning-based ensemble method for single-cell clustering, Sc-GPE [11], is an example of a solution deploying co-association to the problem of ensemble clustering, where a co-association matrix is weighted by the similarity (adjusted rand index) of contributing clustering methods. However, the dependence on a co-association matrix makes this technique intractable for cytometry data. The same limitation applies to SC3 [12], another consensus approach for scRNA-seq employing CSPA for ensemble clustering. Single-cell aggregated (from ensemble) clustering (SAFE-clustering) [13] avoids the need for generating a co-association matrix by applying graph-based methods instead but the implementation only allows a limited number of contributing algorithms to the consensus and is exclusively designed for scRNA-seq.

In contrast to these advances in scRNA-seq data analysis, ensemble clustering methods have yet to be applied to cytometry data. Of note, methods developed for scRNA-seq data analysis may not scale to the size of data encountered in cytometry data analysis, which can be hundreds of times greater. We here benchmarked a range of graph ensemble clustering methods against popular clustering algorithms for cytometry data analysis and present a novel ensemble clustering methodology based on geometric median clustering with weighted voting (GeoWaVe). Compared to graph ensemble clustering methods that have gained popularity in scRNA-seq analysis, GeoWaVe performed favourably on different sets of high-dimensional data generated using cytometry by time of flight (CyTOF) or multicolour flow cytometry. Our findings provide proof of concept for the power of consensus methods to make cytometry data analysis more robust and reproducible.

## 2 Materials and methods

### 2.1 Datasets

Public CytTOF datasets were obtained from open-source repositories [2] and arc-sinh transformed with a standard cofactor of 5. Doublets, debris and dead cells were removed, and ground-truth labels were taken from the original publications, with manual gating by the authors of the *Levine-13* [14], *Levine-32* [14] and *Samusik* [15] datasets. An in-house generated 12-colour flow cytometry dataset was derived from nine patients admitted with acute severe sepsis to the Adult Intensive Care Unit at the University Hospital of Wales in Cardiff, United Kingdom (see **Supplementary Methods**). *Sepsis* data were arc-sinh transformed (standard cofactor of 150) and batch effect corrected using the Harmony algorithm [16]. Each sample was manually gated for single live CD4^+^ and CD8^+^ T cells, Vδ2^+^ γδ T cells and CD161^+^ Vα7.2^+^ mucosal-associated invariant T (MAIT) cells. The identified lymphocyte populations then served as a ground-truth for comparison of results from the clustering algorithms.

### 2.2 Base clustering

The following base clustering algorithms were considered individually, and their outputs served as input to ensemble clustering: FlowSOM (“self-organising map”) [17], PHATE (“potential of heat-diffusion for affinity-based trajectory embedding”) [18] with *k*-means clustering, SPADE (“spanning-tree progression analysis of density-normalized events”) [19], Phenograph [14] and PARC (“phenotyping by accelerated refined community-partitioning”) [20]. These algorithms are computationally efficient and have shown good performance for cytometry data analysis. The output of the base clustering algorithms was used to generate a label-association matrix as input for MCLA, HGPA and HBGF, three graph ensemble clustering algorithms that have been successfully applied to scRNA-seq data [13]. For each base clustering algorithm, experimentation with multiple input parameters was performed to give the best possible performance. The number of clusters generated for each method was determined either as a property of the clustering method (as is the case with Phenograph and PARC), selected from a range of 3-50 using the popular ConsensusClusterPlus method [21], or a fixed value of 20 clusters. The output of the base clustering algorithms was used to generate a label-association matrix (*m* clusters × *n* observations), which served as input for the MCLA, HGPA and HBGF graph ensemble clustering algorithms as implemented using the ClusterEnsembles Python package [22].

### 2.3 Ensemble clustering

For graph ensembles, a required hyperparameter is the number of final partitions in the consensus solution. This problem was addressed in the base clustering algorithms by searching a range of possible clusters and using ConsensusClusterPlus. This approach required sub-sampling the feature space and computing the co-association matrix for each value of *k* (the number of clusters). The cumulative distribution function (CDF) for each co-association matrix was generated, and the optimal *k* chosen where the CDF is maximum. This was applicable to methods such as FlowSOM and SPADE that use a heuristic or down-sampled feature space but was intractable for graph-based consensus clustering techniques that construct graph representations of a *m* × *n* label-association matrix. Therefore, the optimal number of consensus partitions was chosen using internal metrics (metrics that use internal information from the clustering process to evaluate the quality of a clustering, *e*.*g*. the variation within clusters or the degree of overlap between clusters). Ensemble clustering was repeated over a range of *k*; chosen as the smallest and largest number of clusters amongst base clustering algorithms. Four internal metrics, implemented in Scikit-Learn [23], were chosen for their ease of interpretation: Calinski-Harabasz score, Davies-Bouldin index, distortion score and silhouette coefficient. Internal metrics were measured for 1000 events with 100 resamples and the distributions plotted for each *k*. The performance of clustering algorithms in comparison to ensemble methods was evaluated using external metrics that compare cluster results to ground-truth labels, implemented in the Scikit-Learn library [23]: Adjusted Rand Index (ARI), Fowlkes-Mallows Index (FMI) and Adjusted Mutual Information (AMI).

## 3 Results

### 3.1 Graph ensemble clustering methods fail to outperform individual clustering algorithms for cytometry data analysis

Ensemble clustering solutions should take input from informative algorithms suited to the analytical task in question. We here sought to benchmark ensemble methods from the literature using externally and internally generated data, in particular ensemble methods that scale to large cytometry data (greater than 100,000 data points), namely graph-based methods.

Base clustering algorithms and ensemble methods were tasked with clustering three datasets with available ground-truth labels. The *Levine-13* data describe a total of 265,627 bone marrow cells from two healthy human donors and include 13 parameters (**Supplementary Figure S1**) [14]. *Levine-32* describes 167,044 bone marrow cells from a single healthy human donor but at higher resolution with 32 CyTOF parameters (**Supplementary Figure S2**) [14]. Some examples of challenges presented by these two datasets include overlapping monocyte subsets differentiated by CD11b expression in the *Levine-13* data, and small subsets of B-cells differentiated by IgM and IgD expression in the *Levine-32* data. The *Samusik* data describe bone marrow samples with a total of 841,644 cells from 10 C57BL/6J mice and identified 24 populations using 39 CyTOF parameters (**Supplementary Figure S3**) [15]; the branching topology of which offers a unique challenge to any clustering algorithm aiming to partition data in meaningful ways (**Figure 1**). In addition to these three CyTOF datasets, we used an in-house generated 12-colour flow cytometry dataset from sepsis patients (**Supplementary Figure S4**) where the challenge was to identify relatively small and ambiguous populations of unconventional T cells amongst the backdrop of classical CD4^+^ and CD8^+^ T cells (**Figure 1**).

**Figure 1:**
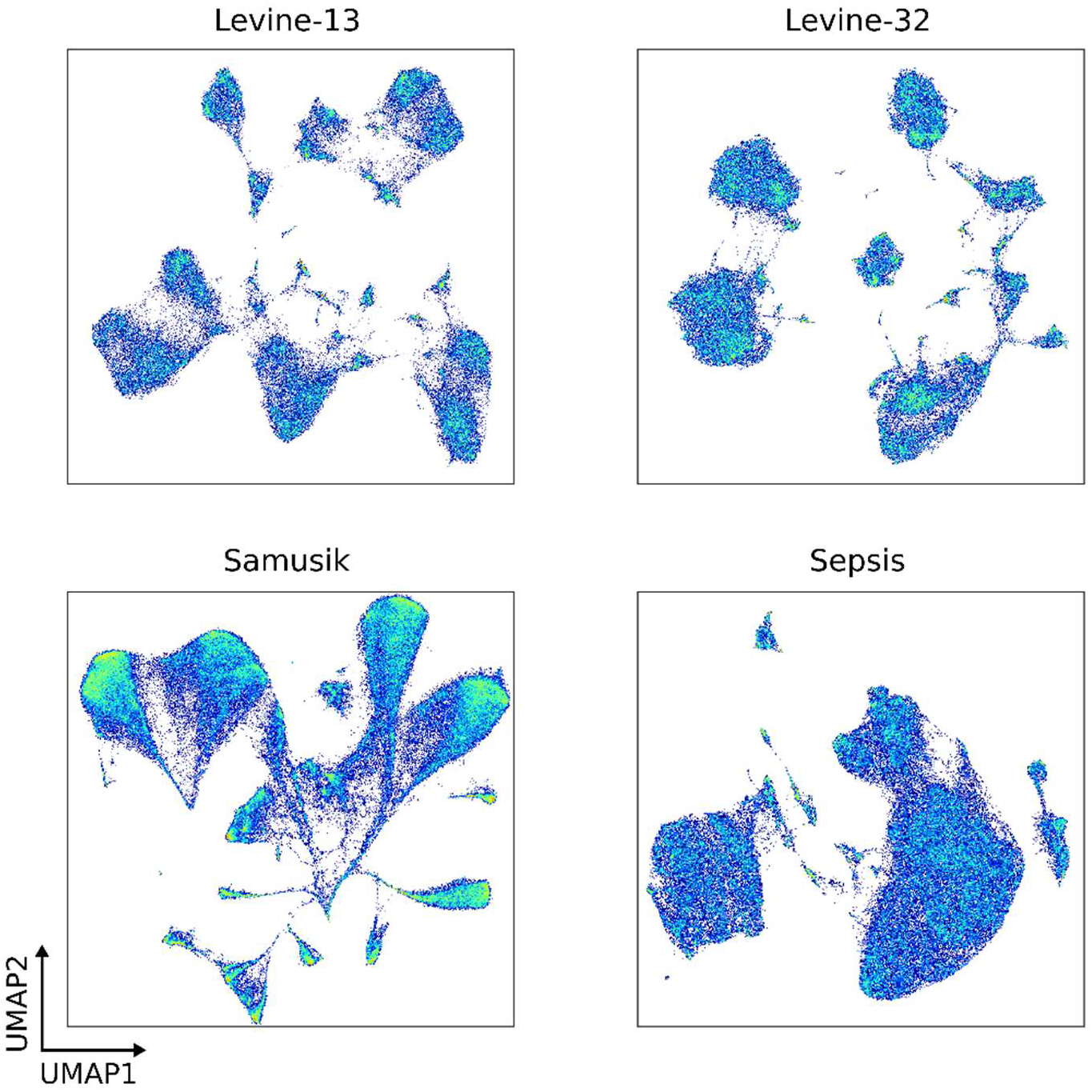
UMAP density plots of the *Levine-13, Levine-32, Samusik* and *Sepsis* data. Colour intensity corresponds to the density of observations in a region of events.

When evaluating the performance of base clustering algorithms in comparison to graph ensemble methods, MCLA offered greater performance compared to the other graph ensemble methods as judged by the adjusted rand index (ARI; **Figure 2**), Fowlkes-Mallows index (FMI; **Supplementary Figure S5**) and adjusted mutual information (AMI; **Supplementary Figure S6**). In the case of the *Levine-13* and *Levine-32* datasets, MCLA provided similar performance to the popular FlowSOM algorithm but overall graph ensemble methods failed to outperform one or more of the base clustering algorithms for all four benchmark datasets.

**Figure 2:**
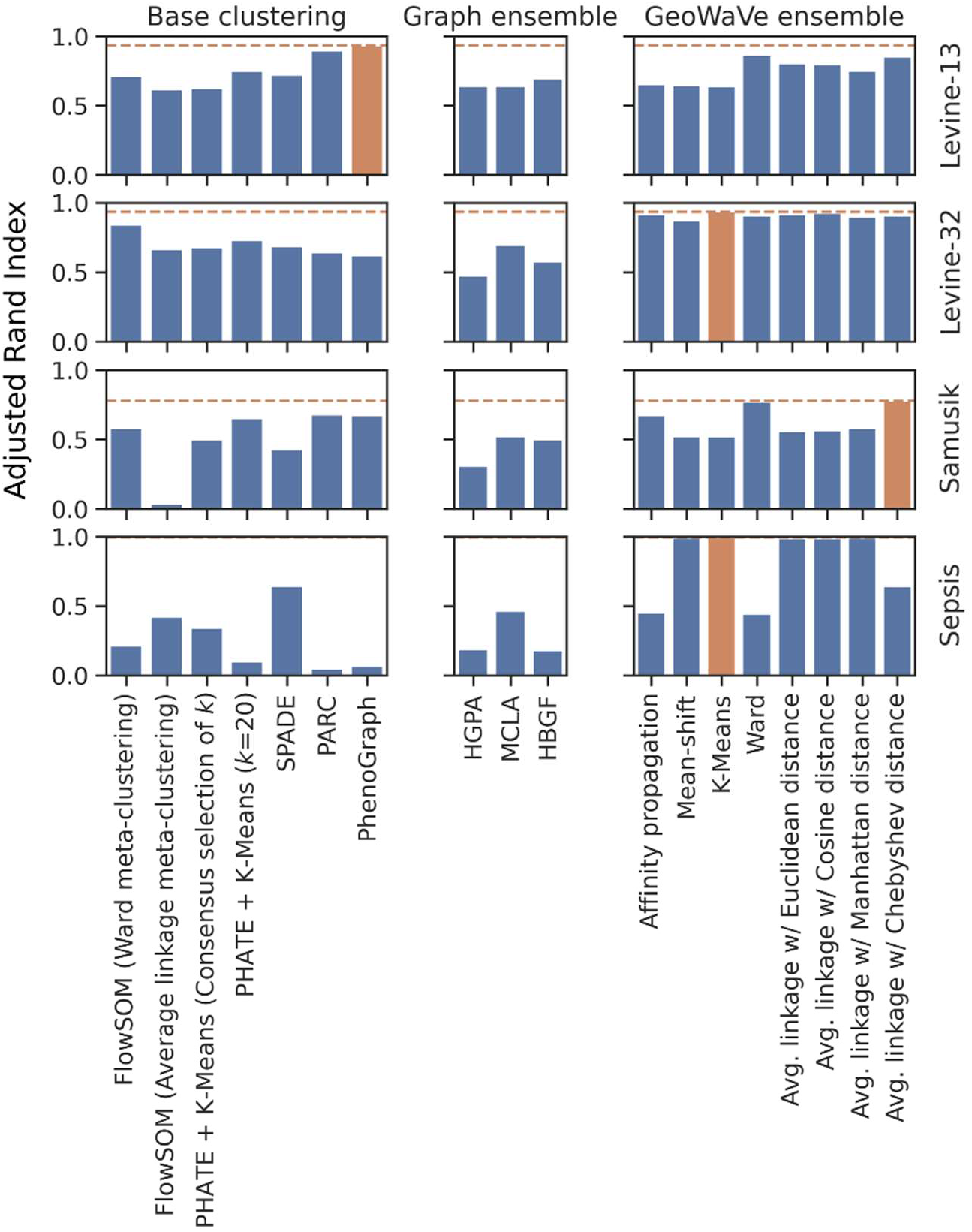
Adjusted rand index (ARI) for base clustering algorithms (left), graph ensemble methods (middle) and GeoWaVe ensemble (right) for the four benchmark datasets. ARI provides a measure of similarity between clusters and ground-truth labels by considering all pairs of observations. Pairs that are assigned to the same or different clusters in the predicted and ground-truth populations are counted and contrasted to mismatched pairs. The Rand Index can be described as a measure of the percentage of correct classifications by the clustering algorithm and is adjusted for chance by estimating the expected rand index using a permutation model, and then normalising by this expectation. ARI scores clustering results between −1 and +1, where random label assignment will be negative or close to zero and perfect clustering will have an ARI close to +1. The best ARI score for each dataset is shown as a dotted orange line, and the best performing method for those data is coloured in orange.

To test whether the performance of graph ensemble methods was a direct result of the method employed for selecting *k*, the number of final consensus clusters (*i*.*e*. the use of internal metrics as shown in **Supplementary Figure S7**), the performance of graph-based clustering algorithms was examined across different values of *k* using external evaluation metrics. HBGF was chosen because it had the best runtime of the three graph ensemble methods (**Supplementary Figure S8**). Here, performance was optimum for low values of *k* despite the number of ground-truth populations being much larger for the *Levine-13* and *Samusik* datasets. The choice of *k* was therefore assumed not to be a factor in the poor performance of graph ensemble methods in this case. Taken together, out findings demonstrate that graph ensemble clustering methods for mass and flow cytometry data performed worse than one or more contributing base clustering solutions.

### 3.2 Geometric median clustering with weighted voting (GeoWaVe): a novel heuristic ensemble clustering algorithm

Graph ensemble methods address issues of computational complexity by deriving the consensus from graph representations of the label-association matrix, rather than from the unmanageable co-association matrix. To improve the performance of consensus approaches, we propose an alternative heuristic ensemble clustering method: geometric median clustering with weighted voting (GeoWaVe), where the clusters generated by base clustering algorithms contributing to an ensemble (such as the examples used in this study, see **Figure 3A**) are summarised by their geometric median (**Figure 3B**). The geometric median (implemented with the hdmedians package [24] and formally defined in **Equation 1**) was chosen over other measures of central tendency because it is robust to outliers, is not necessarily a point from the original data, can handle negative values and is defined in any dimension.

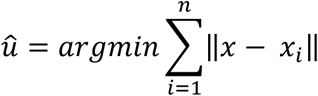

**Figure 3:**
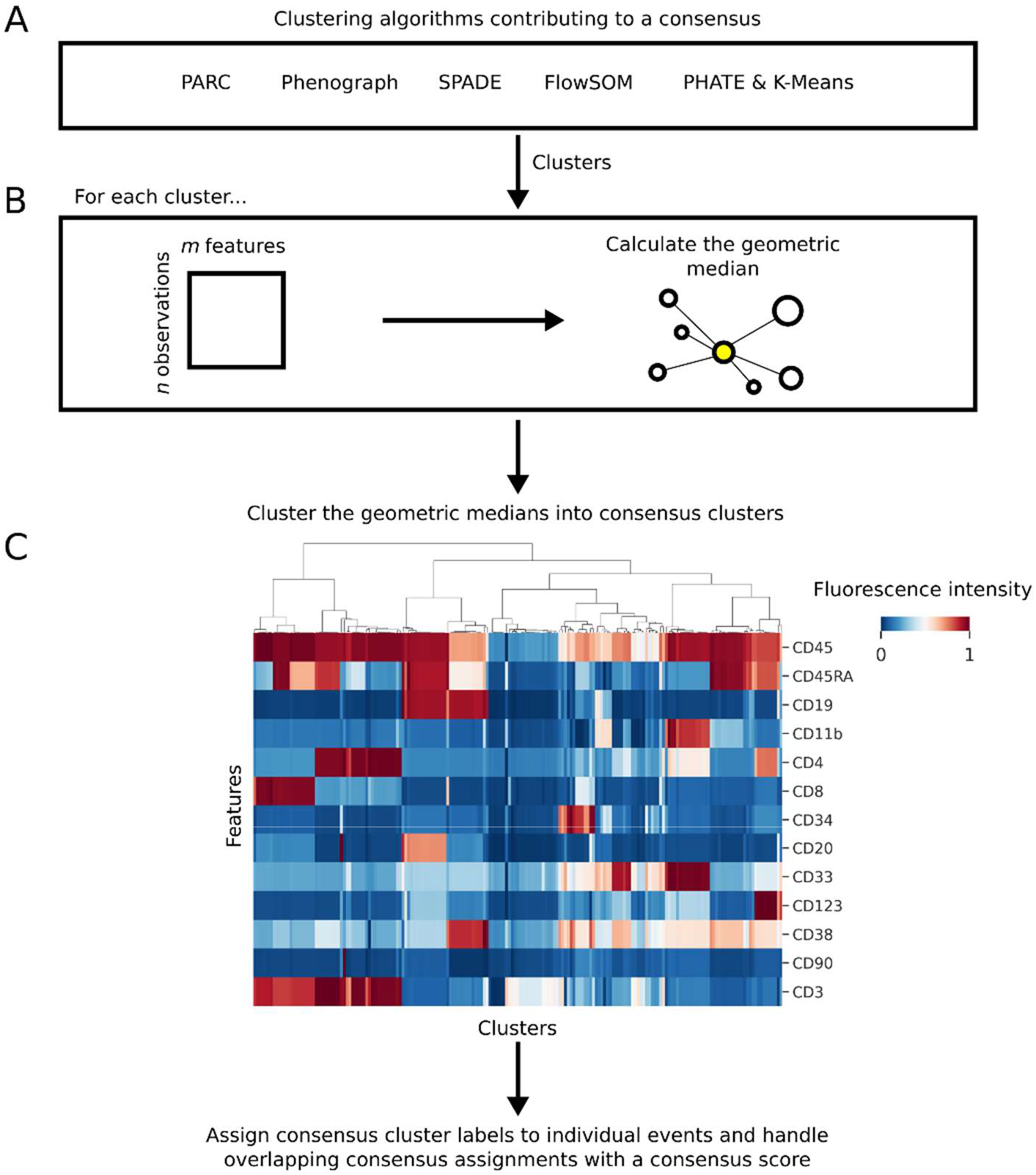
Schematic diagram of the GeoWaVe algorithm. (A) Clusters generated by multiple clustering algorithms are pooled, and (B) the geometric median for each cluster is calculated to create a matrix of *c* clusters. (C) This matrix of cluster geometric medians (clusters of the *Levine-13* data shown here as an example) is clustered into consensus clusters; groups of clusters within similar expression profiles. Consensus cluster labels are then assigned to individual events and overlapping consensus assignments handled with a score that accounts for the distance of the event to the members of each consensus cluster.

Using his approach, a summary of the expression profile of all clusters contributing to the consensus is generated, which can subsequently be clustered into consensus clusters (**Figure 3C**); a consensus cluster being a collection of clusters of similar phenotype. The clusters that contribute to a consensus are overlapping sets, given that each base clustering algorithm is exposed to the same data. Therefore, it is possible that an event can be assigned to more than one consensus cluster. This will occur more frequently for events that sit on the boundary between clusters. To solve this problem, where an event is assigned to multiple consensus clusters a score is calculated for each consensus cluster and the event assigned to the consensus with the maximum score.

The consensus cluster score is calculated as follows: given that a consensus cluster can be defined as a set of clusters *C*, the Manhattan distance between an event *x* and the geometric median (**Equation 1**) of each cluster is computed. The sum of these distances is then divided by the number of clusters within the consensus, given a weighting factor, *p*. This can be expressed formally as **Equation 2:**

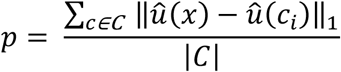

The final score is calculated as the number of clusters within the consensus divided by *p* (**Equation 3**):

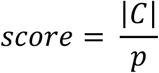

Not all clusters are equally defined, and some may be a poor fit for a given event. To account for this possibility, the majority voting algorithm is weighted by the distance from an event to the centre of each cluster that contributes to a consensus. This method ensures that the consensus an event is assigned to was informed by both the number of supporting algorithms but also the quality of the clusters in that consensus.

The choice of clustering algorithm applied to the geometric medians of clusters is ambiguous in that any number of existing methods may be suitable to the task. The advantage of geometric medians as a heuristic is that the expression profile can be visualised easily as a heatmap (**Figure 3C**) and different clustering methods can be applied and critiqued. This allows the investigator to introduce prior knowledge, such as known phenotypes expected to occur in the data.

### 3.3 Validation of GeoWaVe

To validate GeoWaVe, multiple algorithms for clustering the geometric medians were tried. Affinity propagation and mean-shift were compared because of their ability to select the optimal number of clusters from characteristics of the data. *k*-means and agglomerative hierarchical clustering were also tested, with the optimal number of clusters chosen from a range of clusters using the ConsensusClusterPlus method [21]. For agglomerative hierarchical clustering a variety of linkage methods and distance metrics were tried.

GeoWaVe performance was compared to base clustering algorithms and graph ensemble methods using external evaluation metrics. GeoWaVe outperformed all other methods in three of the four datasets when comparing ARI (**Figure 2**) and FMI (**Supplementary Figure S5**). GeoWaVe also outperformed graph ensemble methods when comparing ARI, FMI and AMI (**Supplementary Figure S6**) but failed to outperform base clustering methods in terms of AMI in the *Levine-13* and *Samusik* data.

The effect of the choice of clustering algorithm applied in GeoWaVe was data specific. For the *Levine-13* and *Levine-32* data, the choice of algorithm was negligible whereas *k*-means and agglomerative hierarchical clustering with average linkage showed a clear advantage in the *Sepsis* data.

### 3.4 GeoWaVe outperforms graph ensemble methods

External evaluation metrics used in the prior section offer performance criteria that are independent of the labels, *i*.*e*. they do not require a like-to-like matching of cluster and ground-truth labels. Instead, measures of similarity between the cluster labels and ground-truth labels were used. Samusik *et al*. [15] and Webber *et al*. [5] alternatively framed such problems in the context of a classification task: a one-to-one mapping of ground-truth labels to clusters was achieved using the Hungarian algorithm such that the sum of F1 scores across ground-truth labels is maximised, and the precision (positive predictive value), recall (sensitivity) and F1 score (harmonic mean of precision and recall) for each ground-truth label are reported.

This procedure was repeated for the clustering algorithms bench-marked in previous sections and the ensemble clustering solutions. **Figure 4** shows the average precision, recall and F1 score for the base clustering algorithms, graph ensemble methods and GeoWaVe with bootstrapped 95% confidence intervals from 1000 rounds of re-sampling. GeoWaVe continued to outperform graph ensemble methods across the four benchmark datasets but failed to match the F1 score obtained by methods such as PhenoGraph and PARC in the *Levine-32* and *Samusik* datasets. While MCLA graph ensemble clustering was more comparable to GeoWaVe in the *Sepsis* data when observing F1 score, GeoWaVe with *k*-means clustering still outperformed MCLA in terms of precision, recall and F1 score.

**Figure 4:**
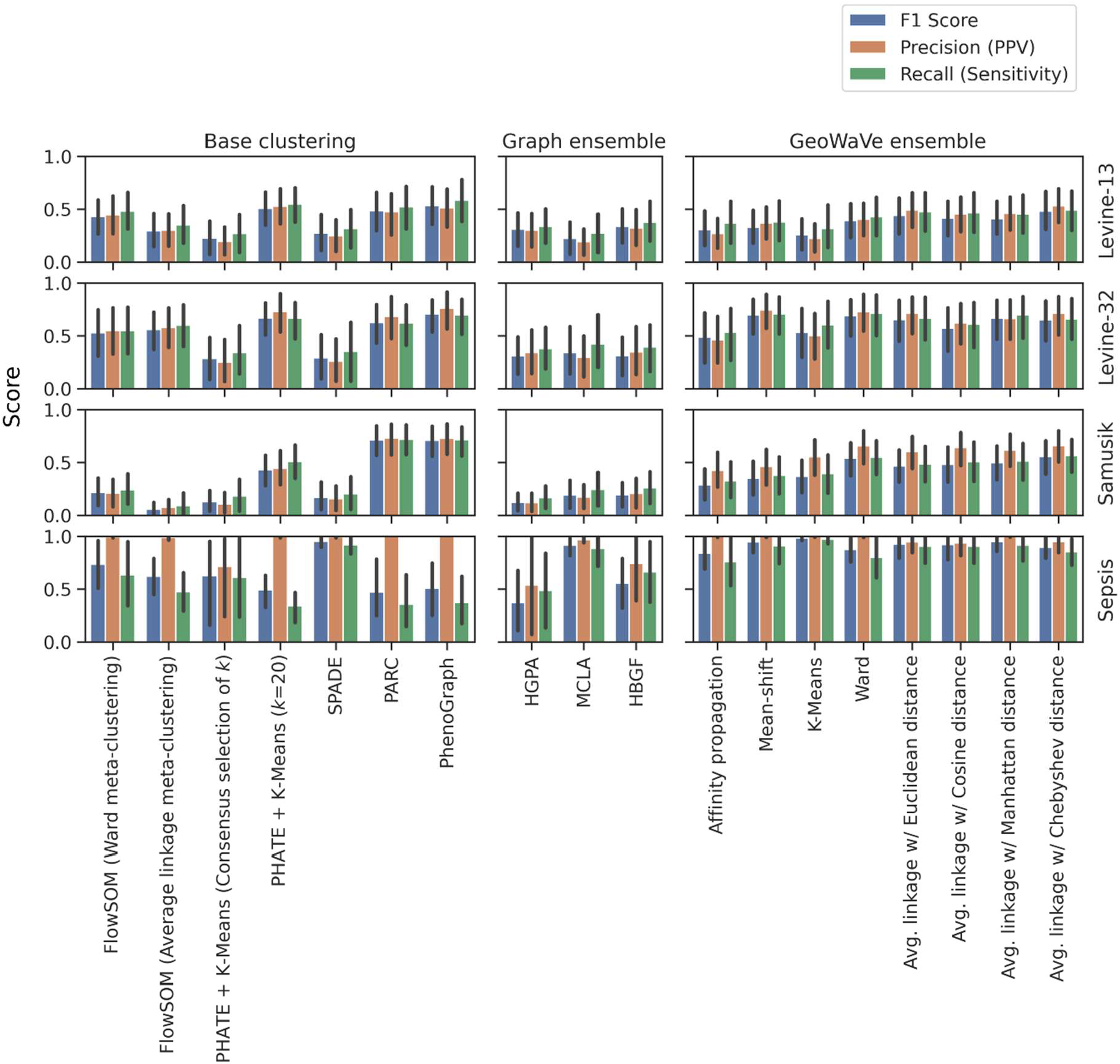
Performance of base clustering algorithms, graph ensembles and GeoWaVe ensembles, after matching cluster labels to ground-truth labels using the Hungarian linear assignment algorithm and maximising the sum of F1 scores across ground-truth label and cluster label pairings. Average precision (positive predictive value; PPV), recall (sensitivity) and F1 score are reported. Error bars represent 95% confidence intervals after bootstrap sampling of population values with 1000 rounds of re-sampling.

An advantage to matching clusters to ground-truth populations using the Hungarian algorithm was the ability to directly compare the performance at the population level. The F1 score for ground-truth populations is shown as a heatmap for the Levine data in **Figure 5** and compares the performance between graph ensemble clustering and GeoWaVe. Each row includes a measure of the population size as an additional heatmap on the y-axis. This approach demonstrated how the superior performance of GeoWaVe compared to graph ensemble methods was a result of improved identification of under-represented populations such as plasma cells and plasmacytoid dendritic cells in the *Levine-13* dataset, and plasma cells, basophils and CD16^+^ natural killer cells in *Levine-32*. The ability of GeoWaVe to outperform graph ensemble methods is supported by observations in the *Samusik* data (**Figure 6**) where multiple populations of moderate size were absent from the output of graph ensemble methods but could successfully be identified by GeoWaVe; examples include natural killer T cells, macrophages and non-classical monocytes. Despite the success of GeoWave in comparison to graph ensemble methods, it still failed to identify very rare subsets in high-dimensional data, for example platelets in the Levine13 dataset and CD34^+^ CD38^+^ CD123^+^ haematopoietic stem and progenitor cells in the *Levine-32* dataset. However, while graph ensemble methods performed better on the *Sepsis* data compared to the other three datasets, minor populations of Vδ2^+^ γδ T cells and MAIT cells were more reliable identified by the GeoWaVe algorithm.

**Figure 5:**
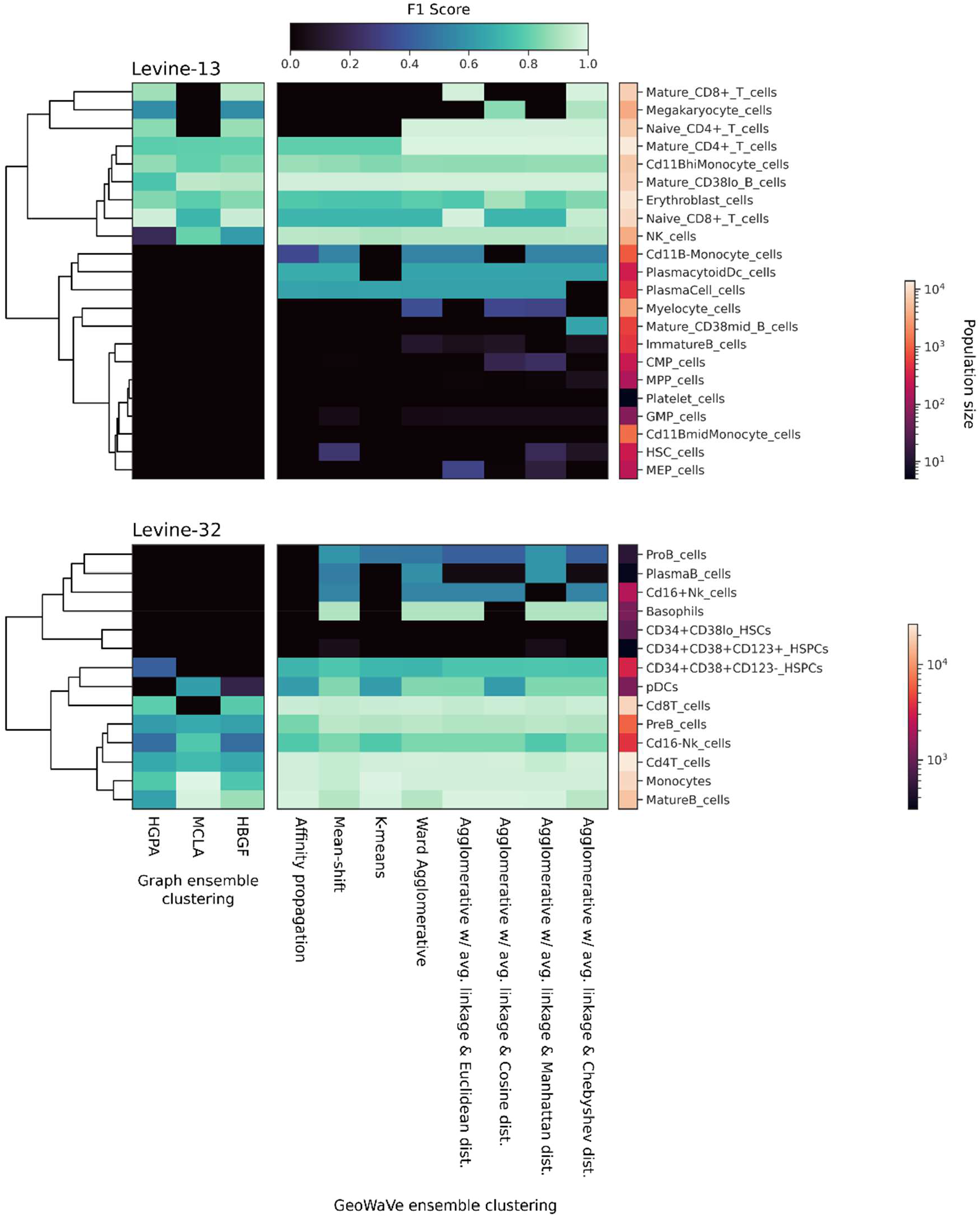
Heatmap of population F1 scores for the *Levine-13* (top) and *Levine-32* (bottom) data, with comparisons between graph ensemble methods (left) to GeoWaVe ensemble methods (right). Ground-truth populations (rows) are coloured by F1 score in the central heatmaps, with darker colours indicating a lower F1 score. On the right y-axis each row is labelled with an additional heatmap that describes the log-normalised size of the population (total number of events).

**Figure 6:**
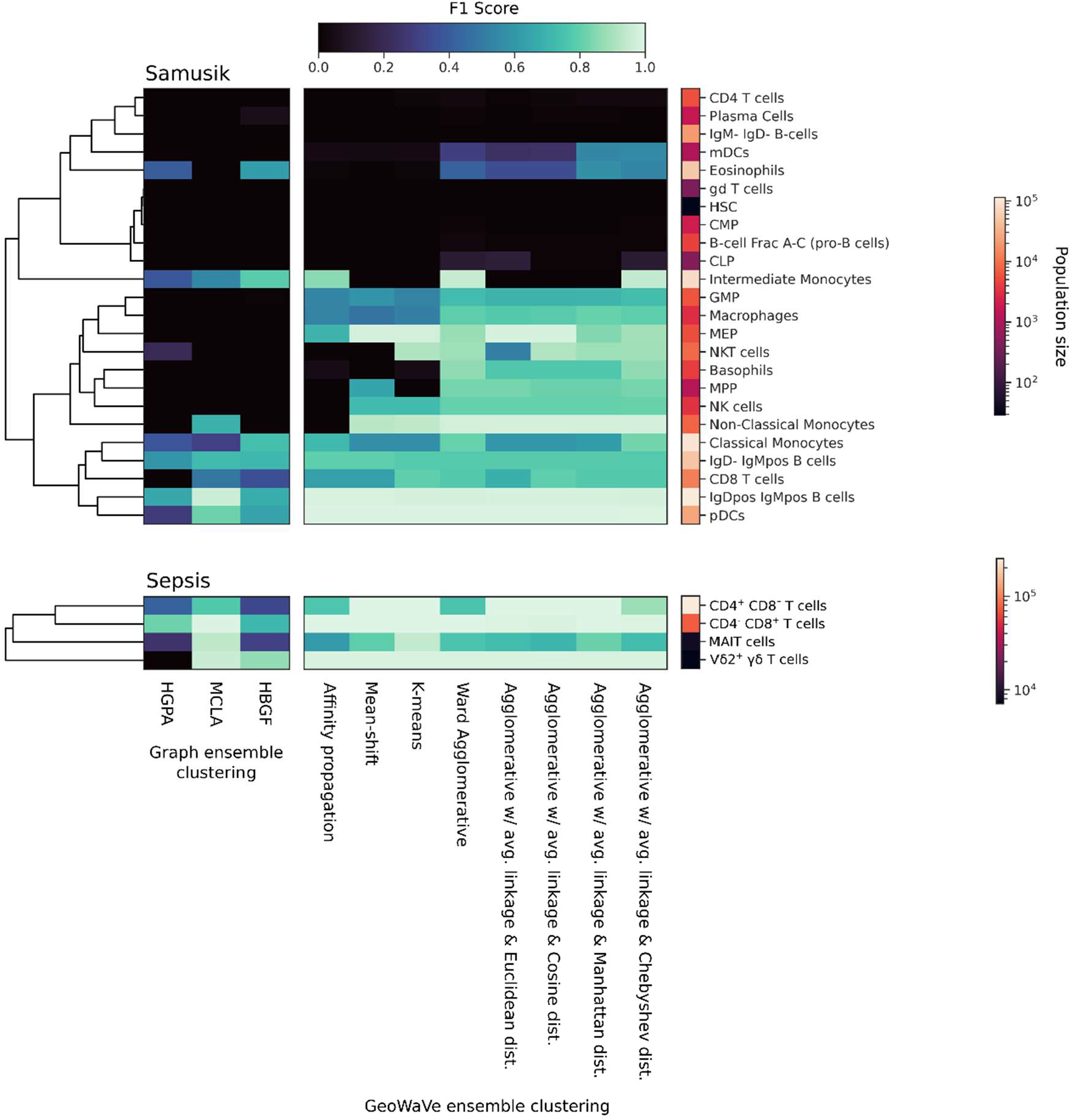
Heatmap of population F1 scores for the *Samusik* (top) and *Sepsis* (bottom) data, with comparisons between graph ensemble methods (left) to GeoWaVe ensemble methods (right). Ground-truth populations (rows) are coloured by F1 score in the central heatmaps, with darker colours indicating a lower F1 score. On the right y-axis each row is labelled with an additional heatmap that describes the log-normalised size of the population (total number of events).

## 4 Discussion

Cytometry has become a cornerstone to biomedical and healthcare research and is widely used in clinical diagnosis. In many pathological conditions, the understanding of disease mechanisms and how to exploit them for patient benefit relies largely on cytometry, including the diagnosis of conditions like leukaemia and HIV infection, and studying antigen-specific responses in vaccine trials. Historically, cytometry data have been processed and analysed manually. Until recently, this was deemed acceptable given that cytometry instruments could only accommodate relatively few parameters in any experiment. Over the past decade, however, the number of available parameters has increased drastically with the advent of multicolour flow cytometry and mass cytometry, allowing characterisation of even minor populations at single cell level and the discovery of novel cell types and new functional features. Traditional approaches no longer suffice – as the number of parameters grows, data analysis is becoming more labour intensive, more subjective and harder to standardise and reproduce across studies and sites. In response to the technological advances, the domain of cytometry bioinformatics is rapidly evolving to provide new computational solutions for data analysis and interpretation such as autonomous gating, supervised classification and unsupervised clustering. Arguably the most impactful technology introduced to this space is dimension reduction for the purpose of data visualisation, such as *t*-distributed stochastic neighbourhood embedding (*t*-SNE), uniform manifold approximation and projection (UMAP) and PHATE. The top clustering algorithms alone have already amassed >12k citations in the scientific literature within a few years and are enabling researchers to make rapid progress in their fields – for instance in the understanding of the immunopathology of COVID-19 that rapidly translated into novel therapies, outcome prediction and vaccine development [25-28].

We here developed GeoWaVe, an ensemble clustering algorithm, as a solution to reduce the variance commonly observed amongst clustering methods in the cytometry literature, where results depend upon hyperparameter choice and the particular context in which they are applied. Presently, there is an absence of a “one size fits all” solution to clustering cytometry data, leaving scientists to rely on exploratory analysis that risks biasing results through data dredging [28]. Ensemble clustering offers an alternative by finding a consensus informed by the results of multiple clustering algorithms exposed to the same data. This multi-view approach theoretically offers robust, consistent and stable solutions [4, 8] without biasing the analysis with the assumptions of a single algorithm. The act of employing ensemble clustering also forces the analyst to compare and contrast the results of multiple algorithms, which can be an informative exercise.

Ensemble clustering presents many challenges that come to bear when applied to complex data such as those generated with cytometry. Unlike supervised classification, there are not a defined number of classes provided by labelled examples. Different algorithms may generate different quantities of clusters, which must be compared and consolidated into consensus clusters. Cytometry data also tends to generate large data that can be difficult to handle with conventional computer resources. This is becoming increasingly relevant for studies that intend on phenotyping hundreds or even thousands of subjects.

An existing ensemble approach that can scale to large data and was included in this study are the graph-based methods, such as HGPA, MCLA and HBGF. These techniques were benchmarked against four independent datasets but failed to outperform individual clustering algorithms such as FlowSOM, PhenoGraph, or SPADE. In response to this, an alternative heuristic ensemble method named GeoWaVe was developed, which was suitable to the nature of cytometry data. Given that the dimensions of cytometry data are not beyond the comprehension of the investigator and meaningful phenotypes can be determined by considering sets of features, we propose to summarise each cluster contributing to a consensus by its geometric median in the feature space. This can for instance be visualised in a heat map. Our study demonstrates that clustering the matrix of these geometric medians can generate informative consensus clusters.

Our analyses showed that GeoWaVe consistently outperformed HGPA, MCLA and HBGF. The use of geometric medians also provided a useful visual aid when choosing the number of consensus clusters to be formed. By visualising the heat map of geometric medians in combination with *t*-SNE, UMAP or PHATE embeddings, a suitable number of partitions can easily be estimated. This allows the investigator to introduce informative priors and select clusters based on knowledge of the underlying biology. If uncertain, a range of partitions can be searched using the ConsensusClusterPlus method [21].

The use of geometric medians as a heuristic is not without limitations. Summarising a cluster using the geometric median tells little of the topology, and a significant loss of information may result in misinformed consensus clusters that are not representative of the data themselves. Additionally, the optimal choice of clustering method applied to the matrix of geometric medians is not immediately apparent and performance can vary depending on the data – for instance, this choice was important to the performance on the *Samusik* dataset, but less relevant for *Levine-32*. Of note, the use of a heuristic means that the run-time of GeoWaVe is fast enough to accommodate hyperparameter tuning. The investigator is therefore encouraged to experiment with different clustering algorithms and hyperparameters and inspect the partitions on the geometric median heat maps and embeddings generated from a suitable dimension reduction technique. Although this fails to remove the exploratory approach to clustering of cytometry data, it introduces the multi-view consensus necessary for robust results.

Webber *et al*. [5] performed a similar assessment of clustering algorithms without the focus on consensus methods and framed their assessment as a classification problem, inspired by the work by Samusik *et al*. [15]. They chose to use F1 score by first mapping clusters to ground-truth labels using the Hungarian algorithm and maximising F1 scores across reference populations. This methodology was repeated in the present study and supported the conclusion that GeoWaVe ensemble methods outperform the graph ensemble methods of HGPA, MCLA and HBGF. Closer inspection of individual population F1 scores revealed that rare cell populations were often not identified by graph ensemble methods. Although identification of these subsets was improved in GeoWaVe, performance was often worse than individual clustering algorithms and some populations, such as platelets in the *Levine-13* data, remained unidentified.

There is a significant flaw in the assessment of clustering performance through F1 score. Mapping clusters to ground-truth labels in such a way implies that a one-to-one relationship must exist between the clusters generated and the reference populations. Clustering analysis can be complicated by sub-structures in data captured as clusters but absent in the ground-truth labels. If the purpose of clustering cytometry data is to identify a precise number of clusters, then this form of evaluation seems justified although one could argue that in such a scenario a supervised classification approach might be more suitable. Clustering analysis tends to be applied in the interest of discovery when the number of clusters is unknown. Despite this flaw it was deemed necessary to replicate the methods of Webber *et al*. [5], which was informative of the role population size plays. It showed that although the consensus clustering of geometric medians outperforms graph-based methods, there is still work to be done to ensure rare cell populations do not go undetected with this technique. It would be advisable that if rare cell populations are suspected to be present, that the consensus is formed by methods with high resolution such as those formed on nearest-neighbour graphs [14, 15, 20].

Future work should focus on more diverse ensemble clustering. In this work, four classes of algorithm were chosen based on their popularity in the cytometry literature and their available implementations. However, there is a wide variety of further clustering algorithms that could be explored for inclusion in ensemble clustering. There are ongoing efforts to address the computational complexity, such as improvements to SC3 [30]. Other solutions to the computational complexity may come from advances in the statistical and computational literature, such as consensus formed on heuristics of cluster similarity using metrics such as the Jaccard index [31]. In the meantime, clustering on geometric medians is likely to be a viable solution for cytometry data analysis. We are confident that the availability of user-friendly but powerful ensemble clustering methods has the potential to represent a major advance in big data analysis, with implications for an improved understanding of cellular responses, clinical conditions, and therapeutic and diagnostic options.

## Acknowledgements

We are grateful to all sepsis patients and their advocates for participating in this study and to the clinicians and nurses for their cooperation. We would also like to thank Jade Cole, Helen Hill, Loïc Raffray and Sarah Baker for their help and advice.

## Data availability statement

The *Levine-13, Levine-32* and *Samusik* datasets are available from Flow Repository, repository number FR-FCM-ZZPH provided by Weber *et al* [2]. The *Sepsis* data underlying this article will be shared on reasonable request to the corresponding author.

