## Supplementary methods, table and figures for "GeoWaVe: Geometric median clustering with weighted voting for ensemble clustering of cytometry data"

*[1]Division of Infection and Immunity, School of Medicine, Cardiff University, Cardiff CF14 4XN, United Kingdom; [2]Adult Critical Care, University Hospital of Wales, Cardiff and Vale University Health Board, Cardiff CF14 4XW, United Kingdom; [3]School of Mathematics, Cardiff University, Cardiff CF24 4AG, United Kingdom; [4]Systems Immunity Research Institute, Cardiff University, Cardiff CF14 4XN, United Kingdom*

[*] To whom correspondence should be addressed

[†] The authors wish it to be known that, in their opinion, the last two authors should be regarded as Joint Senior Authors.

### Supplementary methods: *Sepsis* data

**Ethics statement.** Recruitment of sepsis patients was approved by the Health and Care Research Wales Research Ethics Committee under reference 17/WA/0253, protocol number SPON1609-17 and IRAS project ID 231993 ("Innate-like T cells in sepsis [ILTIS]: Implications for early diagnosis and rescue of immunosuppression") and conducted according to the principles expressed in the Declaration of Helsinki. All participants provided written informed consent for the collection of samples and their subsequent analysis. A waiver of consent system was used when patients were unable to provide prospective informed consent due to the nature of their critical illness or therapeutic sedation at the time of recruitment. In all cases, retrospective informed consent was sought as soon as the patient recovered and regained capacity. In cases where a patient passed away before regaining capacity, the initial consultee's approval would stand.

**Subjects.** Sepsis patients were over 18 years old with a positive diagnosis according to the Third International Consensus Definitions for Sepsis and Septic Shock ('Sepsis-3'). They were cared for in the intensive care unit at the University Hospital of Wales in Cardiff and were recruited within 36 hours of the presumed onset of the infective illness when they already had or would require arterial cannulation as part of standard treatment. Patients were excluded if they were pregnant or breastfeeding, or were females of childbearing age in whom a pregnancy test had not been performed; if they had severe immune deficiency, for example a diagnosis of AIDS or treatment with anti-rejection transplant drugs or high dose corticosteroids; if they had haematologic malignancy or ongoing chemotherapy; if they had severe liver failure (Child's score III or worse); if they were adjudged by the admitting clinician to be unlikely to survive for the duration of the study period regardless of treatment; if they were admitted post-cardiac arrest; or if they had an underlying impairment of higher function that would make it impossible for informed consent to be given upon recovery (*e.g.* severe learning disability). This study cohort comprised a total of $n$=9 sepsis patients, with an age ranging from 18-83 years (median 71 years), 40% of which were female.

**Flow cytometry.** Peripheral blood mononuclear cells were stained after Ficoll-Paque PLUS (Fisher Scientific) separation of blood, using monoclonal antibodies against CD3, CD4, CD8, CD161, TCR-Vα7.2, TCR-Vδ2, TCR-pan-γδ, CD45RA, CCR7 and CD27 (see **Supplementary Table S1**). Cells were acquired on a 16-colour BD LSR Fortessa flow cytometer (BD Biosciences). Live single cells were gated based on side and forward scatter area/height and exclusion of live/dead staining (fixable Aqua; Invitrogen). All data were pre-processed with FlowAI version 3.14 (Bioconductor) to remove poorly acquired events and outliers.

### Supplementary table

| Marker | Conjugate | Clone | Isotype | Company | Dilution |
|---|---|---|---|---|---|
| CD3 | APC/FIRE | SK7 | Mouse IgG1, k | Biolegend | 1:100 |
| CD4 | PE-Cy5.5 | S3.5 | Mouse IgG2α, κ | Life Technologies | 1:200 |
| CD8α | BV711 | RPA-T8 | Mouse IgG1, κ | Biolegend | 1:100 |
| CD14 | V500 | M5E2 | Mouse IgG2α, κ | BD Biosciences | 1:66 |
| CD16 | FITC | 3G8 | Mouse IgG1, κ | BD Biosciences | 1:33 |
| CD19 | V500 | HIB19 | Mouse IgG1, κ | BD Biosciences | 1:66 |
| CD27 | PE-Cy7 | M-T271 | Mouse IgG1, κ | Biolegend | 1:33 |
| CD45RA | PE Dazzle | HI100 | Mouse IgG2b, κ | Biolegend | 1:33 |
| CD57 | FITC | NK-1 | Mouse IgM, κ | BD Biosciences | 1:40 |
| CD161 | APC | 191B8 | Mouse IgG2α, κ | Miltenyi | 1:50 |
| CD197 (CCR7) | BV421 | G043H7 | Mouse IgG2α, κ | Biolegend | 1:25 |
| TCR-pan-γδ | PE-Cy5 | IMMU510 | Mouse IgG1, κ | Beckman Coulter | 1:33 |
| TCR-Vα7.2 | BV605 | 3C10 | Mouse IgG1, κ | Biolegend | 1:33 |
| TCR-Vδ2 | PE | B6 | Mouse IgG1, κ | BD Biosciences | 1:100 |

**Supplementary Table S1:** Flow cytometry staining panel for the analysis of T cells from PBMCs isolated from sepsis patients.

### Supplementary figures

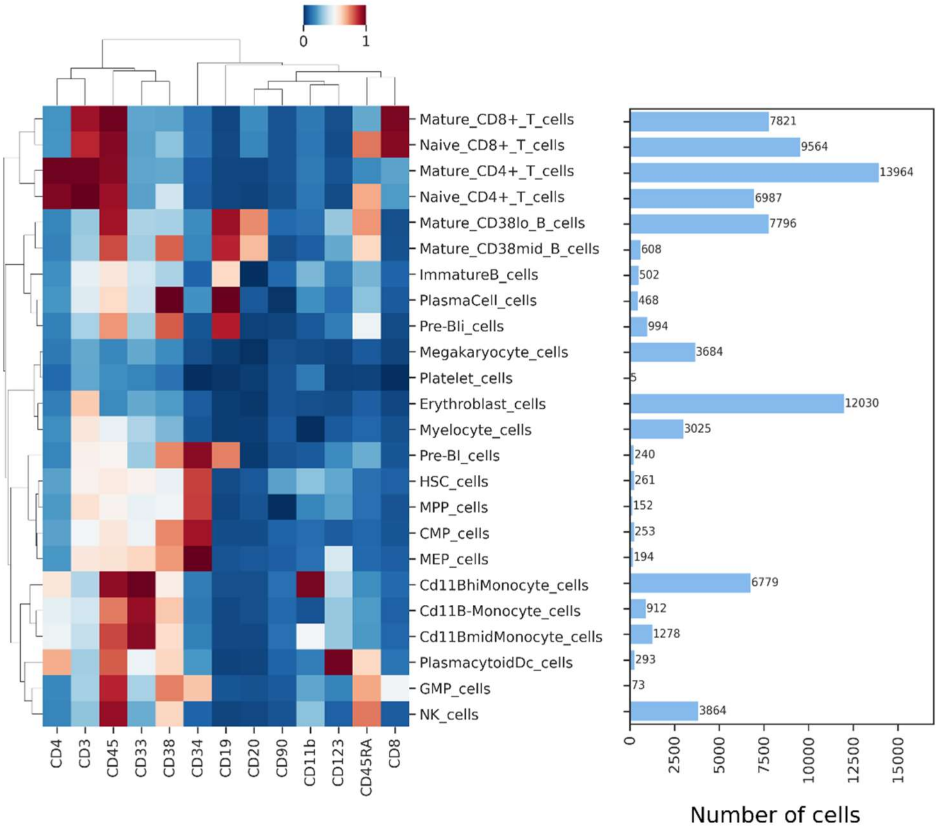


**Supplementary Figure S1:** Expression profile of the 13-parameter *Levine-13* CyTOF data [14] and the total number of observations for each ground-truth population. The heatmap shows the expression intensity of the cell surface markers indicated, normalised to a range between 0 and 1.

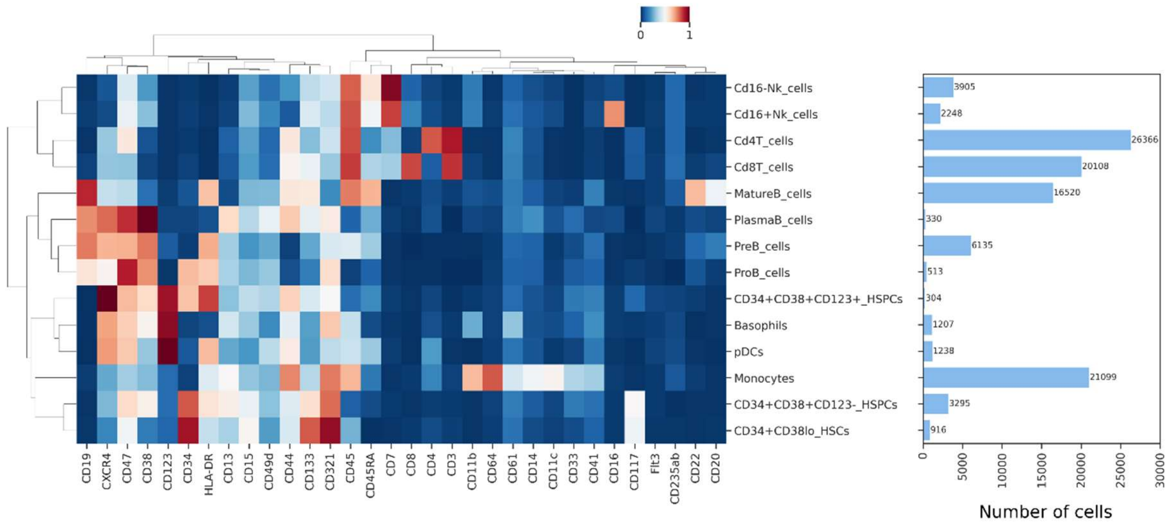


**Supplementary Figure S2:** Expression profile of the 32-parameter *Levine-32* CyTOF data [14] and the total number of observations for each ground-truth population. The heatmap shows the expression intensity of the cell surface markers indicated, normalised to a range between 0 and 1.

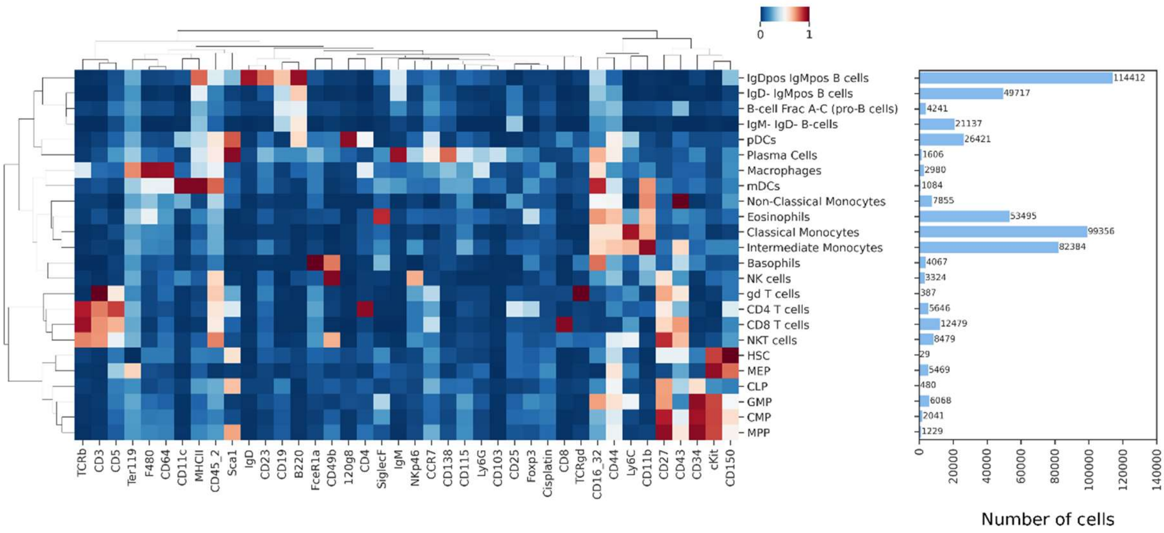


**Supplementary Figure S3:** Expression profile of the 39-parameter *Samusik* CyTOF data [15] and the total number of observations for each ground-truth population. The heatmap shows the expression intensity of the cell surface markers indicated, normalised to a range between 0 and 1.

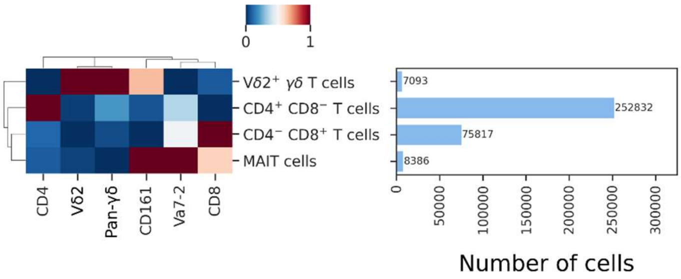


**Supplementary Figure S4:** Expression profile of lineage markers amongst T cell clusters within the 12-parameter *Sepsis* flow cytometry data and the total number of observations for each ground-truth population. The heatmap shows the expression intensity of the cell surface markers indicated, normalised to a range between 0 and 1.

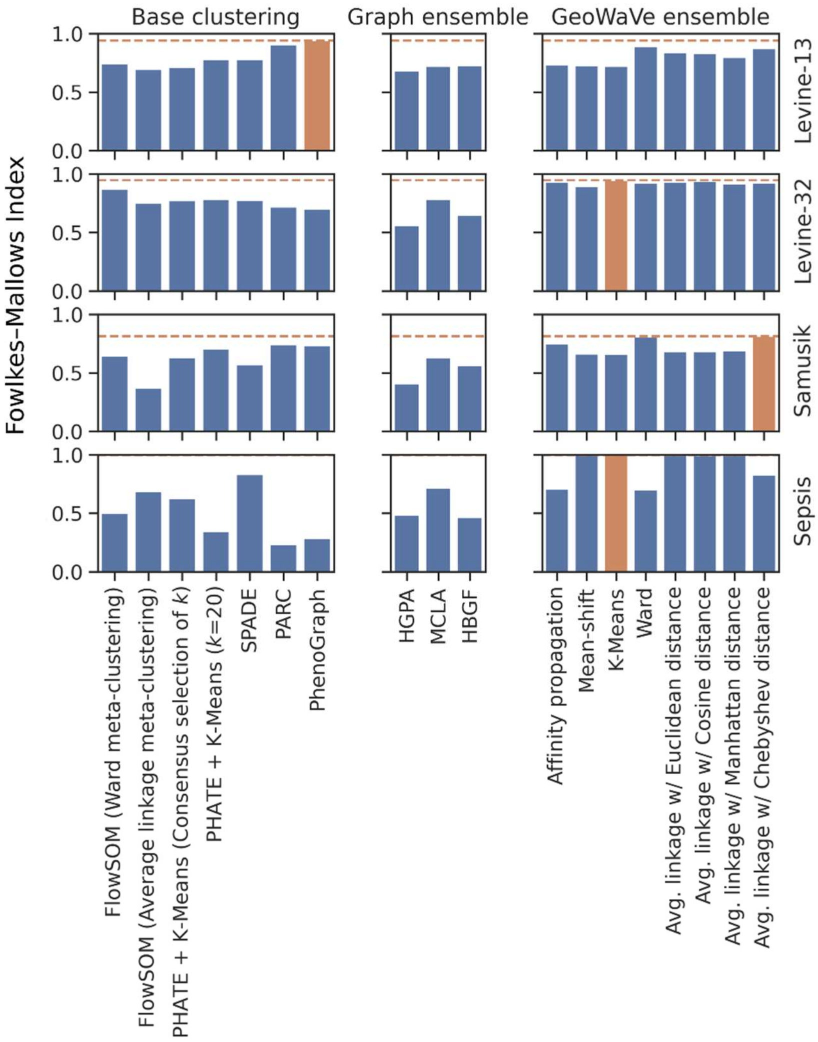


**Supplementary Figure S5:** Fowlkes-Mallows index (FMI) for base clustering algorithms (left), graph ensemble methods (middle) and GeoWaVe ensemble (right) for the four benchmark datasets. FMI is calculated as the square root of the product of pairwise precision (for each ground-truth label, the number of true positives over the number of true positives plus the number of false positives) and pairwise recall (for each ground-truth label, the number of true positives over the number of true positives plus the number of false negatives). FMI is therefore the geometric mean of pairwise precision and recall, has a value between 0 and 1, and a value of 1 will indicate that clusters are identical to the ground-truth labels. The best FMI score for each dataset is shown as a dotted orange line, and the best performing method for those data is coloured in orange.

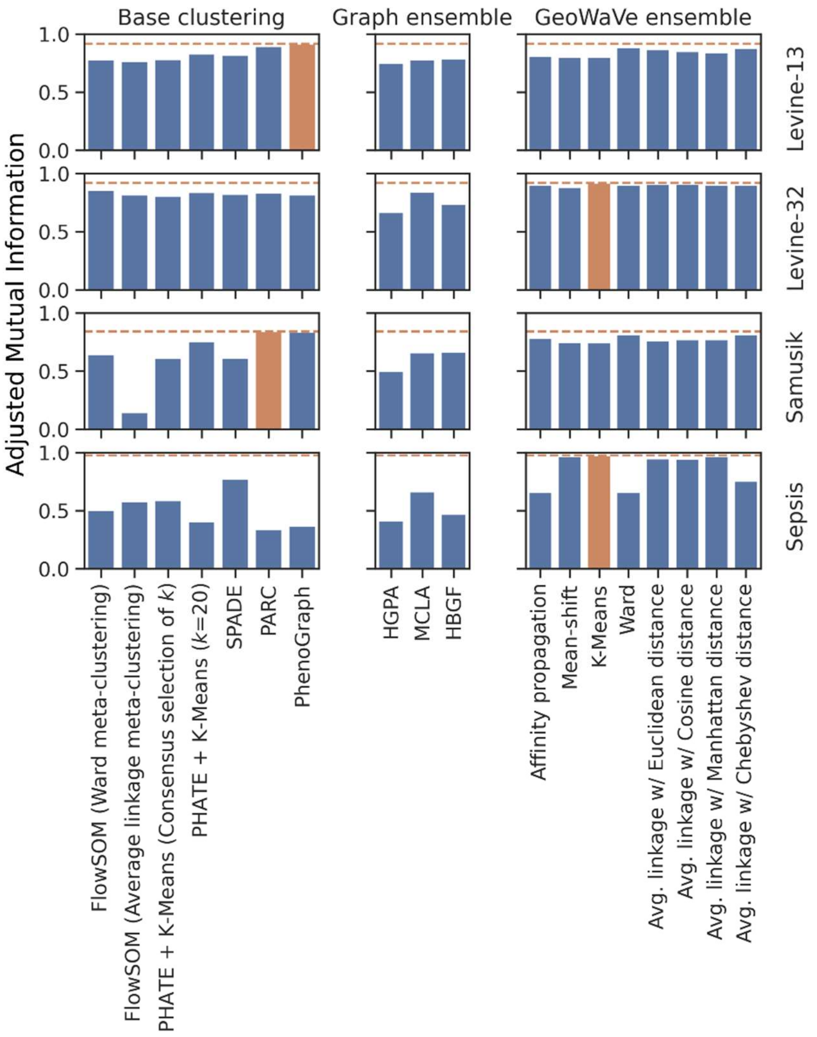


**Supplementary Figure S6:** Adjusted mutual information (AMI) for base clustering algorithms (left), graph ensemble methods (middle) and GeoWaVe ensemble (right) for the four benchmark datasets. AMI is derived from information theory and aims to quantify the amount of shared information between the predicted clusters and the ground-truth populations. Mutual information is not adjusted for chance and will tend to increase as the number of clusters increases, regardless of the quality of additional clusters. To remedy this, AMI first calculates the expected value for mutual information and adjusts for chance in a similar form to the adjusted rand index. AMI scores clustering results between 0 and +1, where random label assignments will give a score of 0 and perfect clustering will have a score of +1. The best AMI score for each dataset is shown as a dotted orange line, and the best performing method for those data is coloured in orange.

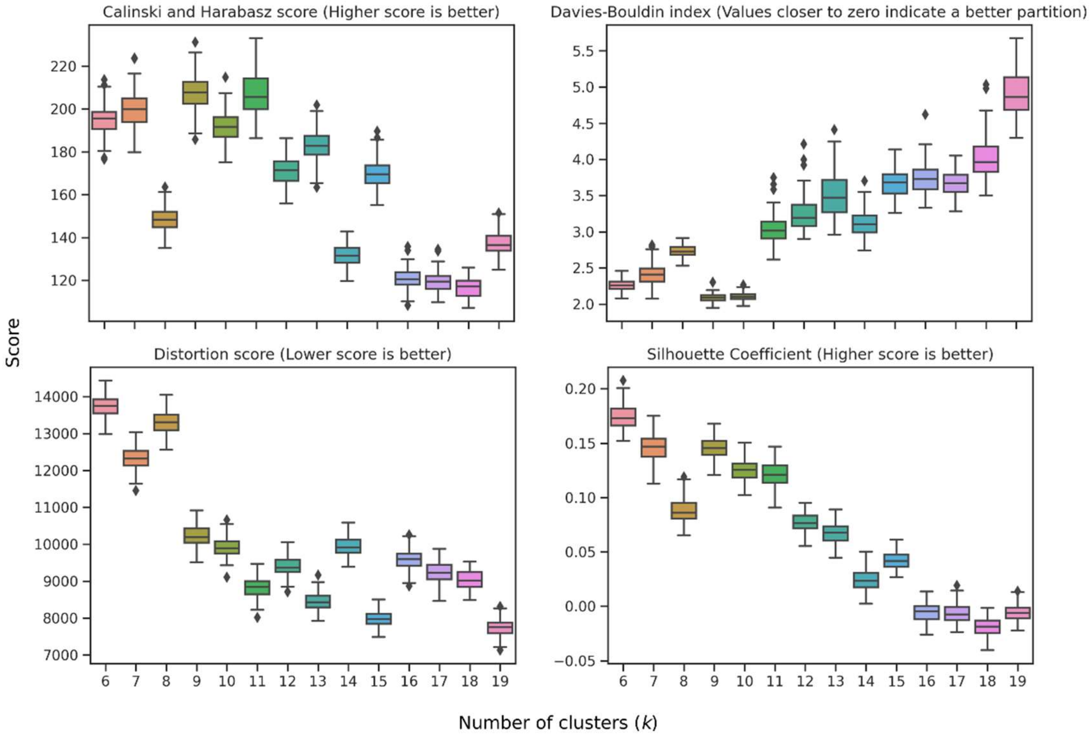


**Supplementary Figure S7:** Internal metrics for a range of final consensus clusters (*k*) as generated by HGPA clustering of *Levine-13* data. The optimal *k* is visually determined as the value where Calinski-Harabasz score and Silhouette coefficient are maximised, whilst Davis-Bouldin index and distortion score are minimised.

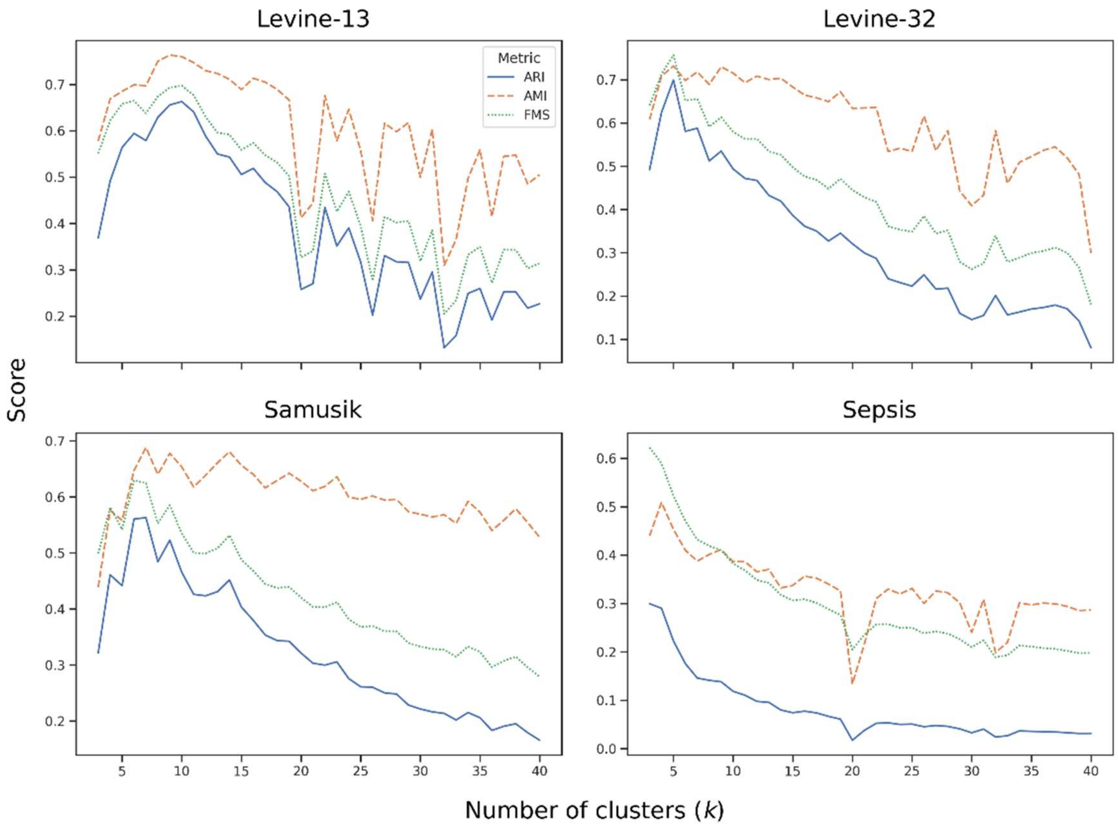


**Supplementary Figure S8:** Adjusted rand index (ARI), adjusted mutual information (AMI) and Fowlkes-Mallows index (FMI) when the number of consensus clusters ($k$) is varied for HBGF ensemble clustering of the *Levine-13* data.
